# PSMA Expression in the Hi-Myc Model; Extended Utility of a Representative Model of Prostate Adenocarcinoma for Biological Insight and as a Drug Discovery Tool

**DOI:** 10.1101/439596

**Authors:** Brian W. Simons, Norman F. Turtle, David H. Ulmert, Diane S. Abou, Daniel L.J. Thorek

## Abstract

Prostate specific membrane antigen (PSMA), also known as glutamate carboxypeptidase II (GCPII), is highly overexpressed in primary and metastatic prostate cancer (PCa). This has led to the development of radiopharmaceuticals for targeted imaging and therapy under current clinical evaluation. Despite this progress, the exact biological role of the protein in prostate cancer development and progression has not been fully elucidated. This is in part because the human PSMA and mouse PSMA (mPSMA) have different patterns of anatomical expression which confound study in the most widely utilized model organisms. Most notably, mPSMA is not expressed in the healthy murine prostate. Here, we reveal that mPSMA is highly upregulated in the prostate and prostate adenocarcinoma in the spontaneous Hi-Myc mouse model, a highly accurate and well characterized mouse model of prostate cancer development. Antibody detection and molecular imaging tools are used to confirm that mPSMA is expressed from early prostatic intraepithelial neoplasia (PIN) through adenocarcinoma.

## Introduction

Targeted imaging and therapeutic delivery to prostate specific membrane antigen (PSMA) is currently of great biomedical interest. Antibodies, peptides and small molecules are all under investigation both preclinically and in late stage clinical trial for diagnostic benefit and therapeutic potential. Indeed, at this time there are 127 active or recruiting clinical trials using PSMA targeted agents or cells for diagnostic or therapeutic purpose in the clinicaltrials.gov database. PSMA protein is expressed in the kidney, gut, brain, peripheral nerves, salivary glands and prostate. In malignant tissues, expression can be detected in the neovasculature of multiple cancer types, and is highly upregulated in prostate cancer tissues. Expression in prostate cancer increases with disease stage of and the majority of both soft-tissue and osseous metastases highly express the antigen (*1*,*2*).

Discovered independently in several different tissues including as glutamate carboxypeptidase II (GCPII) in the gut, PSMA in the prostate and N-acetyl-L-aspartyl-L-glutamate peptidase I of NAAG peptidase I (NAALADase I) in the brain, the enzyme encoded by *FOLH1* has different functions in different tissues. Within the brain it catabolizes NAAG, the most prevalent peptidic neurotransmitter, to N-acetylaspartate and glutamate; in the intestine it cleaves terminal glutamates from dietary folic-polyglutamates. The biological role of PSMA in the prostate and in cancer is not fully known.

Mice are commonly utilized to more deeply understand the etiological basis and impact of protein expression, as well as to evaluate imaging agents and experimental agents in models of disease, and should be a useful tool to investigate PSMA function. Human and mouse PSMA share 86% sequence identity, with identical enzymatic activity (*3*). However, PSMA expression patterns in mice are different than those found in man. Some reports have indicated low (*3*) or a lack (*4*) of expression of mouse FOLH1 (*Folh1*) in the mouse prostate, while others report moderate to high expression in normal prostate and in the TRAMP genetically engineered mouse model (GEMM) of prostate cancer (*5*–*7*). Investigation of mPSMA in cancer progression and development have used transgenic mPSMA expressing tumor models (*8*) and PSMA knockout mice crossed with TRAMP mice (*7*,*9*,*10*) to suggest that PSMA promotes prostate cancer formation and progression. Some of this discrepancy may be due to multiple similar family members that can interfere with specific detection of PSMA. Here we use multiple PSMA specific detection methods, including high-sensitivity non-invasive *in vivo* positron emission tomographic (PET) imaging, to show little or no PSMA expression in normal mouse prostate, but highly upregulated expression in the Hi-Myc model of prostate cancer. This unexpected finding has significant import to improve our understanding of PSMA in disease biology and provides a validated research model for drug evaluation.

## Results

In order to confirm the ability of antibodies used for immunohistochemistry to specifically detect PSMA expression in mouse tissues, we performed protein detection on tissues known to express PSMA. In kidney, PSMA expression was detected in proximal renal tubules and in Bowman’s capsule with the expected membranous staining pattern (Figure 1A). Specificity was confirmed by repeating the staining conditions in PSMA knockout mice (Figure 1B). PSMA expression was undetectable in all examined tissues from knockout mice, suggesting no apparent cross reactivity with other PSMA family members. Little to no expression was detected in normal prostate (Figure 1C) from wild type mice.

**Figure 1.**
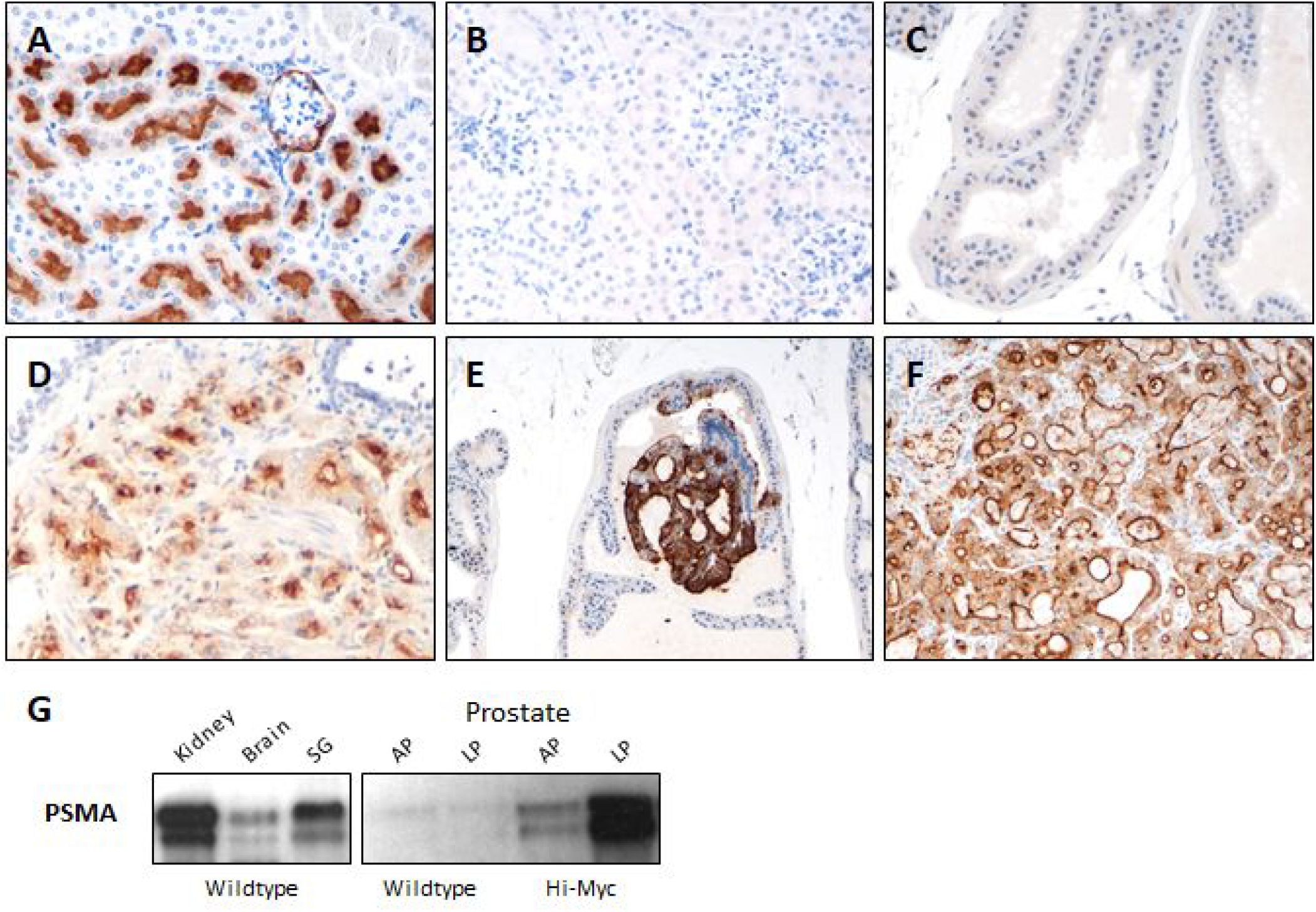
Specific Detection of PSMA in the Hi-Myc Model of Prostate Cancer. Immunohistochemistry using anti-PSMA antibodies in wild type mouse kidney (A) shows high expression in proximal tubules. Negative staining in the kidney of a *Folh1* knockout mouse (B) confirms antibody specificity. Normal mouse prostate (C) shows no staining. Human prostate cancer (D) shows high PSMA expression concentrated at cell membranes. Hi-Myc PIN (E) and adenocarcinoma (F) show high PSMA expression with a similar pattern. Western blot (G) shows similar results with expected expression in kidney, brain, and salivary gland (SG), but not in wild type prostate anterior lobe (AP) or lateral lobe (LP). Hi-Myc prostates show PSMA expression proportional to the expected tumor burden in AP (low burden) and LP (high burden).

In human prostate cancer samples (Figure 1D), high levels of PSMA expression using the same antibody was detected and revealed the expected pattern of membranous staining with concentration at the luminal border of epithelial cells. Surprisingly, in Hi-Myc mice (Figure 1E-F), PSMA expression was detected in PIN lesions (Figure 1E) and in invasive adenocarcinoma (Figure 1F) with a membranous expression pattern similar to human cancer. Tissue lysates prepared from wild type and Hi-Myc mice were similarly assayed for PSMA expression using the same antibody used for immunohistochemistry (Figure 1G). A doublet band of the expected size (approximately 100 kDa) was detected in kidney, brain, and salivary gland. A faint band was present in wild type prostate, and moderate expression was present in the anterior prostate, with high expression present in the lateral prostate lobe of Hi-Myc mice. Myc expression and cancer burden in the Hi-Myc mice is not equal in all prostate lobes but following the previously reported pattern of AR-expression, with very high Myc expression and high volume disease present in the lateral prostate lobe, and scattered Myc expression and low volume disease present in the anterior prostate (*11*).

High levels of expression were detected in preinvasive (PIN) lesions and invasive adenocarcinoma, so we assayed for PSMA expression at very early time points to determine when PSMA expression began in Hi-Myc mice. In this model, the AR-driven Probasin promoter drives Myc expression as early as 2 weeks of age, and PIN lesions begin to develop shortly after Myc expression begins. At 4 weeks of age, no PSMA was detectable despite the presence of Myc-induced PIN lesions in the prostate (Figure 2A). At 8 weeks of age, PIN is widespread in the Hi-Myc prostate, and PSMA is expressed at high levels in PIN lesions (Figure 2B). PSMA expression continues at 6 months (Figure 2C), and most mice will have PSMA positive invasive lesions at this time point.

**Figure 2.**
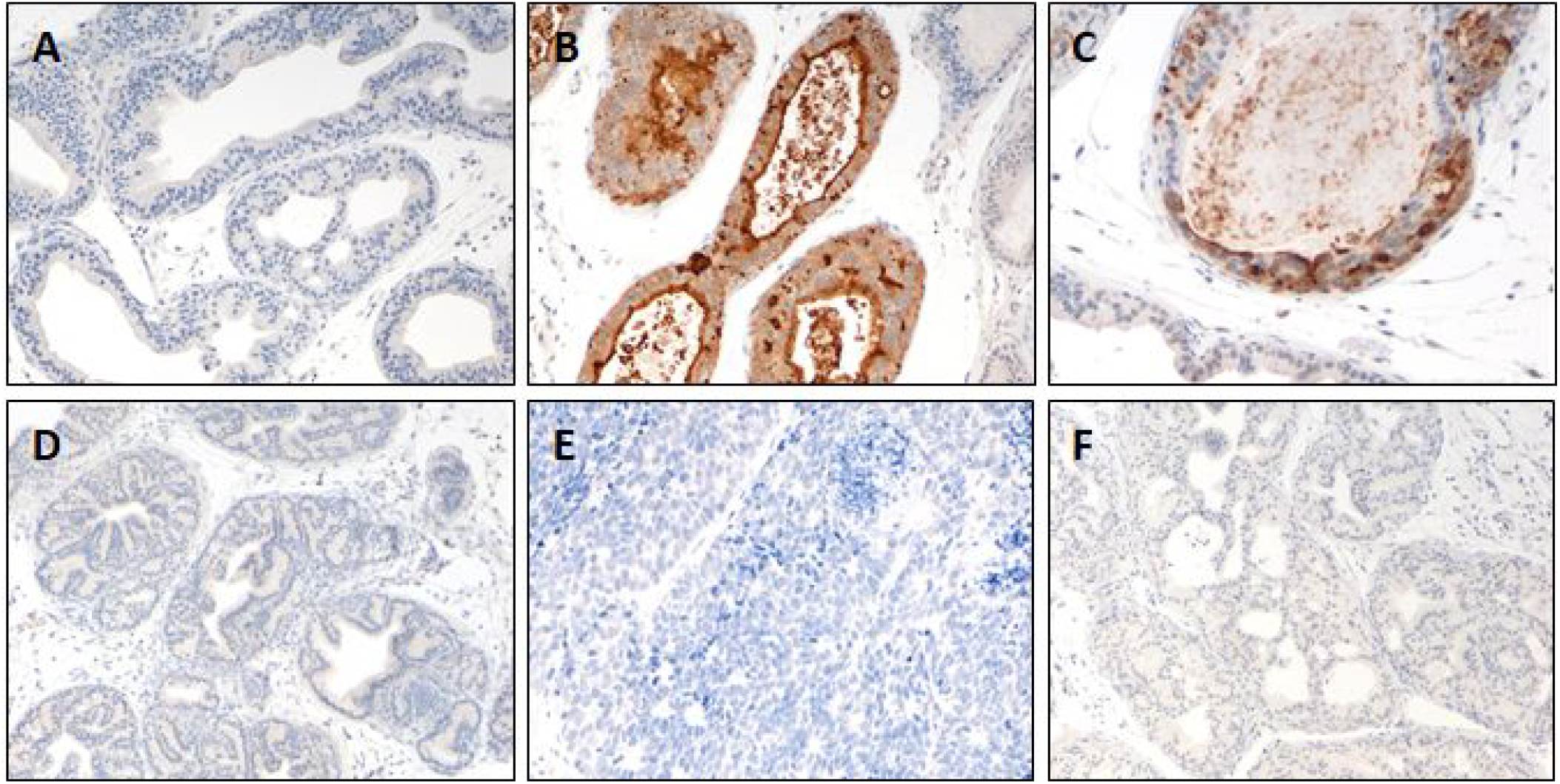
PSMA Expression is Limited to the Hi-Myc Model. Hi-Myc mice (A-C) show no PSMA expression in PIN at 4 weeks of age (A), but high expression is present at 8 weeks (B) and 6 months (C). TRAMP PIN (D), TRAMP invasive tumors (E), and PTEN deleted tumors (F) show no PSMA expression.

Myc expression and PIN development in Hi-Myc mice is heterogeneous in most prostate lobes, and areas of normal prostate cells interspersed between PIN lesions served as negative internal controls. In order to determine if PSMA expression is a feature of all prostate cancer models, or Hi-Myc in particular, we assayed for PSMA expression in TRAMP (Figure 2D-E) and prostate specific PTEN deleted mice (Probasin-Cre x PTEN fl/fl, Figure 2F). No PSMA expression was detected in TRAMP PIN lesions (Figure 2D), invasive TRAMP lesions (Figure 2E), or PB-Cre PTEN lesions (Figure 2F).

To this point, all detection of PSMA expression was accomplished using a single antibody reagent in genetically engineered control and diseased mice. To further confirm our findings of PSMA expression in a mouse model, and to accomplish this in an *in vivo* setting, we utilized the exemplary GCPII-ligand 2-(3-(1-carboxy-5-[(6-[^18^F]fluoro-pyridine-3-carbonyl)- amino]-pentyl)-ureido)-pentanedioic acid ([^18^F]-DCFPyL) (*12*,*13*). Animals under isoflurane anesthesia were administered 9.25 MBq [^18^F]-DCFPyL and imaged at 60 minutes post-injection. Representative volume rendered datasets are shown in Figure 1. PET volumes are shown with dual intensity scales (for without and with bladder segmentation) as bladder activity concentration exceeds 100%IA/mL. In the two perspectives one can visualize the caudally oriented dorsal and lateral prostate lobes, with some signal ascending rostrally along the anterior lobular projections of the prostate (also known as the coagulating glands). No prostatic focus of [^18^F]-DCFPyL-PET signal is detected in the TRAMP mice (notably with significantly greater volume of disease in these age matched animals (*14*). Likewise, wild type mice have no detectable prostatic signal (dissection results shown below).

To confirm these *in vivo* results, the genitourinary package was promptly removed in its entirety, microdissected at 4° C and arrayed (Figure 4A). The tissues were rescanned and uptake, corrected for decay and injection activity, was analyzed (Figure 4B,C). Recapitulating the *in vivo* imaging and pathobiological findings, signal in the lobes of the prostate bearing Myc-driven disease displayed considerable accumulation of [^18^F]-DCFPyL.

## Discussion

In these experiments, we have established the presence of PSMA in a spontaneous, autochthonous, and well regarded model of prostate adenocarcinoma development. Beginning with immunohistochemical evaluation, we have verified that the reagents used to interrogate mouse and human tissues were specific for mouse and human PSMA. Knockout animals, which have intact *GcpIII* (*Folh2*), did not stain positive and served to underline the specificity of our approach. In contrast with some previous reports, we detected minimal to no expression of PSMA in normal and TRAMP prostates. We relied on detection of PSMA protein using a monoclonal antibody and a small molecule, while some reports have detected *Folh1* RNA. The reason for this discrepancy is unclear, but differing sensitivity of these methods or detection of highly similar homologs may explain these differences.

There are many agents under development for improved detection and targeted therapy of prostate adenocarcinoma (*15*–*19*). The widespread use of high-sensitivity and robust methods to detect PSMA in particular in the locally advanced and metastatic settings by PET and SPECT are poised to revolutionize management for those afflicted with the disease. Further, many of these agents can be derivatized to deliver radioisotopes with therapeutic intent (*20*–*22*). This creates the urgent need to better understand the underlying biology of the target protein, its expression in disease states and its response to novel and conventional therapies. We pursued *in vivo* imaging in order to evaluate whether the expression levels detected by molecular biology and histological tools were sufficient to annotate disease status non-invasively in the Hi-Myc model. Utilizing the urea-based [^18^F]-DCFPyL, a high-resolution GCPII-ligand with optimized pharmacokinetics in late-stage clinical evaluation (NCT03181867, NCT02981368), expression in the prostate can be detected in these animals. This is notable given the considerable background signal provided by the bladder when using this renally excreted agent (Figure 3). No detectable signal was observed in wild-type or TRAMP model animals. Excised tissue imaging was performed in order to more clearly visualize expression in the lobes of the prostate. Differential disease development is known in this model; within the dorsolateral and ventral lobes are concentrations of proliferating lesions at earlier time points and to a greater degree (*11*,*15*,*23*) and to quantitate uptake at the sub-organ level. These values also correspond with immunohistochemical findings in Figure 1.

**Figure 3.**
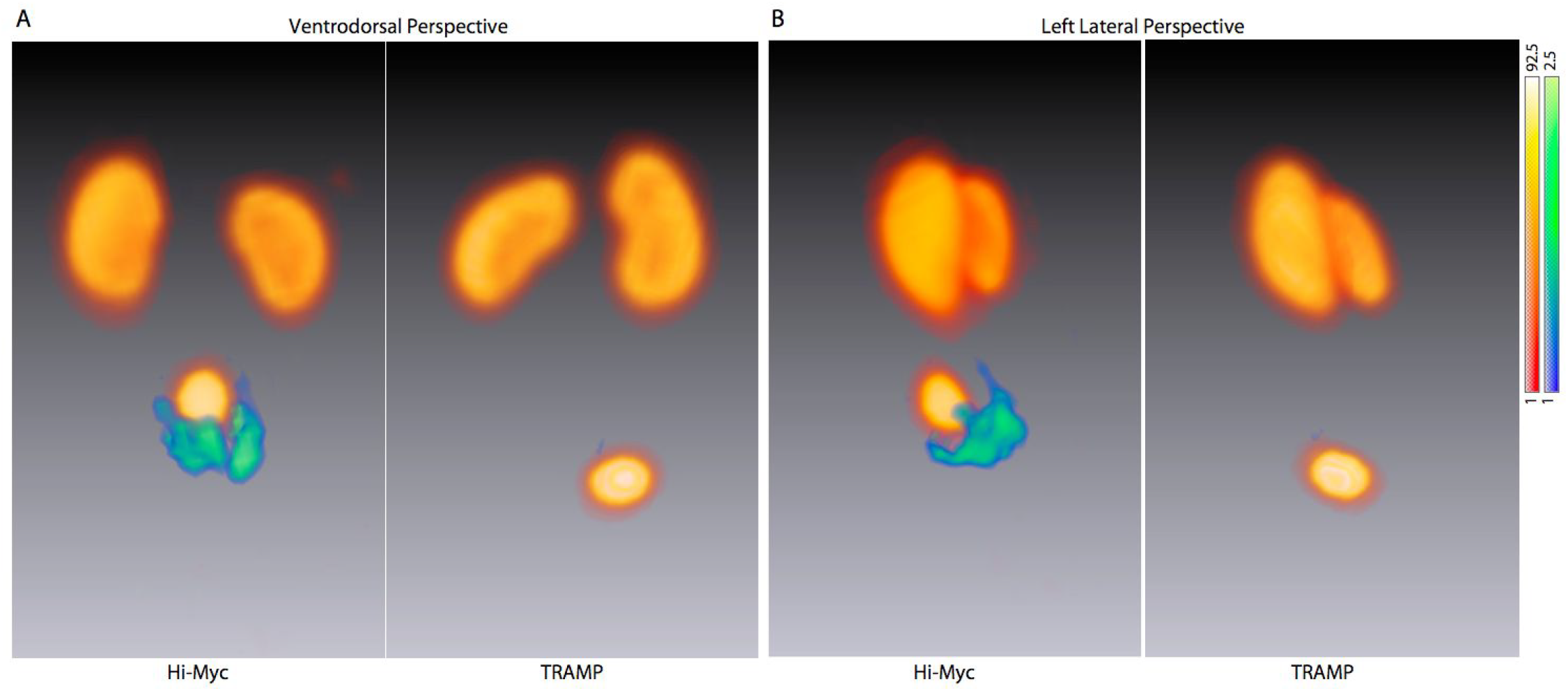
*In Vivo* Imaging of Prostatic PSMA Expression. Three dimensional rendered [^18^F]DCFPyL-PET images of representative age-matched Hi-Myc (FVB) and TRAMP tumor-bearing mice. (A) The ventral to dorsal perspective are shown, and (B) the corresponding left lateral volume projections are shown. These volume rendered images are generated by a weighted average of intensities along the projection axis of the given perspective, and are thus semiquantitative. A red-yellow scale has been used to show intense uptake and clearance of [^18^F]DCFPyL through the kidneys and bladder; a blue-green scale was then used on the thresholded prostate. The region of contrast adjacent the bladder in the Hi-Myc mice corresponds with the morphology of the rodent prostate; no corresponding region was seen in TRAMP tumor-bearing mice. Confirmatory *ex vivo* analyses are shown in Figure 4.

**Figure 4.**
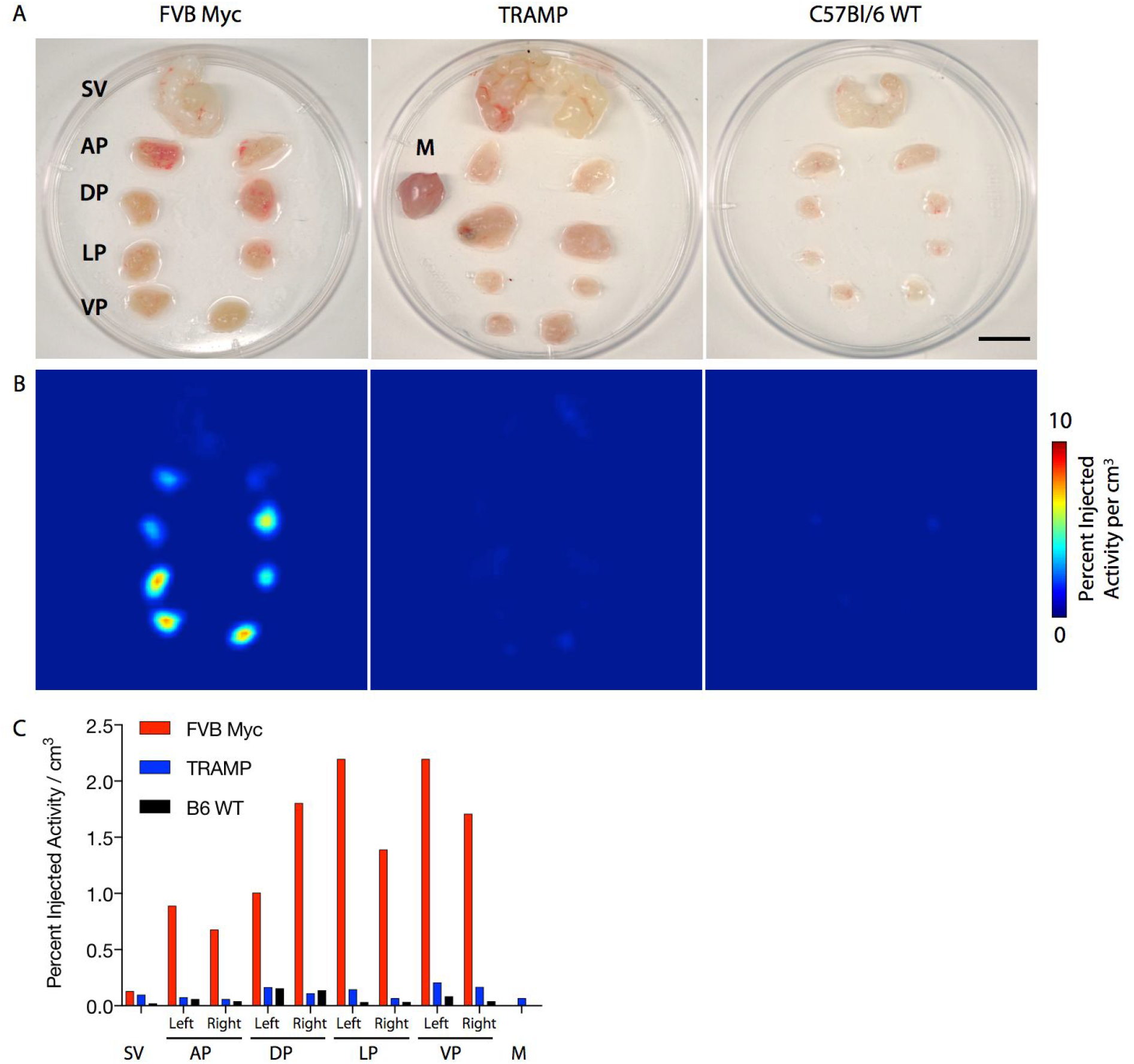
PSMA-Radioligand Uptake in Excised and Dissected Prostate. A) Photograph of representative dissected genitourinary package including Seminal Vesicle (SV) and Anterior Prostate (AP), Dorsal (DP), Lateral (LP) and Ventral (VP) from three mice strains Hi-Myc^FVB^, TRAMP and WT-C57Bl/6. Muscle tissue for reference background is included in the central sample (scale bar 5 mm). B) *Ex vivo* microPET acquisitions of the dissected tissues as above. Uptake is present in the Hi-Myc samples, notably in the VP, and DLP. C) Quantification of volumes of interest of the tissues recapitulates the *in vivo* and *ex vivo* imaging results.

There is no definitive explanation for the absence of GCPII expression in normal mouse prostate, in contrast to that in man. One partial explanation for this is that models of prostatic disease are limited by several factors, key among these is that rodents, used for the bulk of biological evaluation in mammalian systems, do not naturally develop prostate cancer. Another is that within mice, PSMA expression patterns vary from sites in man. Some sites, including the gut, kidney and brain share common GCPII profiles across species. For example, while it has become evident that GCPII is often expressed in the neovasculature of several types of cancer in man, the presence of the protein is absent in xenograft models of cancer in mice (*24*).

Our finding of PSMA expression in the pre- and cancerous prostate of a well regarded mouse model is significant as it may be useful in elucidating the reason for the differences in expression in human and mouse tissues. Moreover, the use of PSMA-targeting agents is now being firmly established in clinical practice for diagnostic and therapeutic intent. This model and the ability to direct such ligands to diseased tissues to recapitulate clinical absorbed dose profiles at the tumor and off-target tissues is of considerable value to assess and optimize methods. To conclude, our work extends the biological utility of this genetically engineered model system for use towards understanding the role of this abundant prostate cancer cell-surface marker in disease progression, and in drug development applications for anti-PSMA imaging and therapy evaluation.

## Methods

### Mice

All experimental procedures were approved by the Johns Hopkins Institutional Animal Care and Use Committee (IACUC). Wild type C57BL/6J, FVB/NJ, and TRAMP mice (C57BL/6-Tg(TRAMP)8247Ng/J) were obtained from Jackson Laboratories (Bar Harbor, ME, Stocks 664, 1800, and 3135). FVB-Tg(ARR2/Pbsn-MYC)7Key (Hi-Myc, Strain 01XF5) mice were obtained from NCI Mouse Repository (Frederick, MD). PSMA-null mice were a gift of Dr. Joseph Neale (Georgetown University, Washington, DC, USA), who had received them originally from Warren Heston (Cleveland Clinic Lerner Research Institute, Cleveland, Ohio, USA) (*25*). Briefly, this GEMM has excised exons 3-5 of *GcpII/Folh1* and backcrossed in C57Bl/6 >10 generations to generate homozygous PSMA-null mice. PSMA is undetectable in the knockout by PCR or western blot, and knockout mice express no basal phenotypic differences (*26*,*27*). Probasin-Cre x PTEN^fl/fl^ mice were generated as previously described (*28*). Genotyping was performed using primer sets and protocols recommended by the vendor. Genomic DNA for PCR was isolated from tails.

### Immunohistochemistry

Slides were deparaffinized and rehydrated before steaming in EDTA pH 8 (Invitrogen) for 40 minutes. Endogenous peroxidases were quenched with BLOXALL (Vector Labs), and the slides were blocked for one hour with Serum Free Protein Block (Dako). Slides were incubated with antibodies directed against PSMA (Cell Signaling Technology rabbit monoclonal antibody (D7I8E). Staining was visualized with ImmPRESS Polymer detection kit and ImmPACT DAB (Vector Labs).

### Western blot

Lysates were prepared from flash frozen tissue samples using RIPA buffer with protease and phosphatase inhibitors added (Sigma). After separation by SDS-PAGE, samples were transferred to PVDF membranes and blocked with 5% NFDM solution. Blots were incubated with antibodies directed against PSMA (Cell Signaling Technology rabbit monoclonal antibody (D7I8E) diluted in 5% NFDM for 1 hour, followed by peroxidase labelled secondary antibody and visualized with SuperSignal chemiluminescent substrate (Thermo Fisher).

### *In vivo* and *vivo* PET

Age matched wild-type (FVB and C57Bl/6), TRAMP, Hi-MYC^FVB^ and Hi-MYC^C57Bl/6^ male mice were administered 9.25-13 MBq (250-300 μCi) [^18^F]DCFPyL intravenously (n=2). After 1 h uptake time, animals were anesthetized with isoflurane and placed prone on in the center of the field of view of a dedicated rodent microPET R4 (Concorde Microsystems). List-mode data were captured in order to acquire 25×10^6^ coincident events (approximately 15 minutes) using a γ-ray energy window of 350–750 keV, and a coincidence timing window of 6 ns. Following imaging, representative animals were sacrificed and the genitourinary (GU) package and bladder removed. The excised GU was placed in a petri dish on an ice slurry and dissected using a stereomicroscope (Zeiss Stemi 2000). The separated tissues were placed in a clean 4 cm diameter dish with 0.5 mL of ice water and rescanned in the microPET R4 for whichever transpired first: 5×10^6^ coincident events or 30 minutes. Data were sorted into 2-dimensional histograms by Fourier re-binning, and images were reconstructed by a maximum *a posteriori* and 3D filtered back projection. Image data were normalized to correct for non-uniformity of response of the PET, dead-time count losses, positron branching ratio, and physical decay to the time of injection and quantification determined relative to injected activity (to yield percent injected activity per gram (%IA/g) or per mL (%IA/mL)). No attenuation, scatter, or partial-volume averaging correction was applied. Images were analyzed by using ASIPro (Concorde Microsystems). Regions of interest were delineated on sequential two dimensional slices to compute uptake values per mL of tissue specimen. For *in vivo* 3D rendered PET data a weighted average of the intensities along projections through the tomographic image is computed, and overlaid with a second volume in which the bladder and kidneys have been thresholded out. The image intensity scales represent the full image intensity space of 0-100% (white-red) and 0-2.5% (blue-green) using Amira 6.5 (FEI). [^18^F]DCFPyL was obtained from the Johns Hopkins University School of Medicine PET Center and synthesized with high specific activity (>150GBq/μmol) as previously reported (*29*).

## Acknowledgements

Funding for this work was supported in part by R01 CA201035 (DLJT). We thank the members of the Johns Hopkins University PET Center under the leadership of Dr. Robert F. Dannals, and Dr. Martin G. Pomper of the Division of Nuclear Medicine and Molecular Imaging in the Department of Radiology and Radiological Science for [^18^F]DCFPyL. Assistance, material and support were also provided by Dr. Karen Sfanos, Corey Porter, Dr. Barbara Slusher, and Dr. Diane Peters (JHU). The authors also respectfully acknowledge Dr. Don Coffey.

## Bibliography

1. Bravaccini S, Puccetti M, Bocchini M, et al. PSMA expression: a potential ally for the pathologist in prostate cancer diagnosis. Sci Rep. 2018;8:4254.

2. Kasperzyk JL, Finn SP, Flavin R, et al. Prostate-specific membrane antigen protein expression in tumor tissue and risk of lethal prostate cancer. Cancer Epidemiol Biomarkers Prev. 2013;22:2354–2363.

3. Bacich DJ, Pinto JT, Tong WP, Heston WD. Cloning, expression, genomic localization, and enzymatic activities of the mouse homolog of prostate-specific membrane antigen/NAALADase/folate hydrolase. Mamm Genome. 2001;12:117–123.

4. Aggarwal S, Ricklis RM, Williams SA, Denmeade SR. Comparative study of PSMA expression in the prostate of mouse, dog, monkey, and human. Prostate. 2006;66:903–910.

5. Schmittgen TD, Zakrajsek BA, Hill RE, et al. Expression pattern of mouse homolog of prostate-specific membrane antigen (FOLH1) in the transgenic adenocarcinoma of the mouse prostate model. Prostate. 2003;55:308–316.

6. Yang D, Holt GE, Velders MP, Kwon ED, Kast WM. Murine six-transmembrane epithelial antigen of the prostate, prostate stem cell antigen, and prostate-specific membrane antigen: prostate-specific cell-surface antigens highly expressed in prostate cancer of transgenic adenocarcinoma mouse prostate mice. Cancer Res. 2001;61:5857–5860.

7. Caromile LA, Dortche K, Rahman MM, et al. PSMA redirects cell survival signaling from the MAPK to the PI3K-AKT pathways to promote the progression of prostate cancer. Sci Signal. 2017;10.

8. Yao V, Parwani A, Maier C, Heston WD, Bacich DJ. Moderate expression of prostate-specific membrane antigen, a tissue differentiation antigen and folate hydrolase, facilitates prostate carcinogenesis. Cancer Res. 2008;68:9070–9077.

9. Greenberg NM. Transgenic models for prostate cancer research. Urol Oncol. 1996;2:119–122.

10. Gingrich JR, Greenberg NM. A transgenic mouse prostate cancer model. Toxicol Pathol. 1996;24:502–504.

11. Iwata T, Schultz D, Hicks J, et al. MYC overexpression induces prostatic intraepithelial neoplasia and loss of Nkx3.1 in mouse luminal epithelial cells. PLoS One. 2010;5:e9427.

12. Szabo Z, Mena E, Rowe SP, et al. Initial Evaluation of [18F]DCFPyL for Prostate-Specific Membrane Antigen (PSMA)-Targeted PET Imaging of Prostate Cancer. Mol Imaging Biol. 2015;17:565–574.

13. Chen Y, Pullambhatla M, Foss CA, et al. 2-(3-{1-Carboxy-5-[(6-[18F]fluoro-pyridine-3-carbonyl)-amino]-pentyl}-ureido)-pentanedioic acid, [18F]DCFPyL, a PSMA-based PET imaging agent for prostate cancer. Clin Cancer Res. 2011;17:7645–7653.

14. Kaplan-Lefko PJ, Chen T-M, Ittmann MM, et al. Pathobiology of autochthonous prostate cancer in a pre-clinical transgenic mouse model. Prostate. 2003;55:219–237.

15. Thorek DLJ, Watson PA, Lee S-G, et al. Internalization of secreted antigen-targeted antibodies by the neonatal Fc receptor for precision imaging of the androgen receptor axis. Sci Transl Med. 2016;8:367ra167.

16. McDevitt MR, Thorek DLJ, Hashimoto T, et al. Feed-forward alpha particle radiotherapy ablates androgen receptor-addicted prostate cancer. Nat Commun. 2018;9:1629.

17. Zhang H, Desai P, Koike Y, et al. Dual-Modality Imaging of Prostate Cancer with a Fluorescent and Radiogallium-Labeled Gastrin-Releasing Peptide Receptor Antagonist. J Nucl Med. 2017;58:29–35.

18. Zhang H, Chen J, Waldherr C, et al. Synthesis and evaluation of bombesin derivatives on the basis of pan-bombesin peptides labeled with indium-111, lutetium-177, and yttrium-90 for targeting bombesin receptor-expressing tumors. Cancer Res. 2004;64:6707–6715.

19. Vilhelmsson-Timmermand O, Santos E, Thorek DLJ, et al. Radiolabeled antibodies in prostate cancer: a case study showing the effect of host immunity on antibody bio-distribution. Nucl Med Biol. 2015;42:375–380.

20. Barinka C, Rojas C, Slusher B, Pomper M. Glutamate Carboxypeptidase II in Diagnosis and Treatment of Neurologic Disorders and Prostate Cancer. Curr Med Chem. 2012;19:856–870.

21. McDevitt MR, Ma D, Lai LT, et al. Tumor therapy with targeted atomic nanogenerators. Science. 2001;294:1537–1540.

22. Kratochwil C, Bruchertseifer F, Giesel FL, et al. 225Ac-PSMA-617 for PSMA-Targeted α-Radiation Therapy of Metastatic Castration-Resistant Prostate Cancer. J Nucl Med. 2016;57:1941–1944.

23. Ellwood-Yen K, Graeber TG, Wongvipat J, et al. Myc-driven murine prostate cancer shares molecular features with human prostate tumors. Cancer Cell. 2003;4:223–238.

24. Nguyen DP, Xiong PL, Liu H, et al. Induction of PSMA and Internalization of an Anti-PSMA mAb in the Vascular Compartment. Mol Cancer Res. 2016;14:1045–1053.

25. Bacich DJ, Ramadan E, O’Keefe DS, et al. Deletion of the glutamate carboxypeptidase II gene in mice reveals a second enzyme activity that hydrolyzes N-acetylaspartylglutamate. J Neurochem. 2002;83:20–29.

26. Rais R, Jiang W, Zhai H, et al. FOLH1/GCPII is elevated in IBD patients, and its inhibition ameliorates murine IBD abnormalities. JCI Insight. 2016;1.

27. Gao Y, Xu S, Cui Z, et al. Mice lacking glutamate carboxypeptidase II develop normally, but are less susceptible to traumatic brain injury. J Neurochem. 2015;134:340–353.

28. Wang S, Gao J, Lei Q, et al. Prostate-specific deletion of the murine Pten tumor suppressor gene leads to metastatic prostate cancer. Cancer Cell. 2003;4:209–221.

29. Ravert HT, Holt DP, Chen Y, et al. An improved synthesis of the radiolabeled prostate-specific membrane antigen inhibitor, [(18) F]DCFPyL. J Labelled Comp Radiopharm. 2016;59:439–450.

